# An Early Devonian permineralized rhyniopsid from the Battery Point Formation of Gaspé (Canada)

**DOI:** 10.1101/200246

**Authors:** Kelly C. Pfeiler, Alexandru M.F. Tomescu

## Abstract

The Emsian deposits of the Battery Point Formation (GaspéCanada) host the most diverse Early Devonian flora in North America. While most of this diversity has been described from plant compressions, the permineralized component of the flora is incompletely explored. Based on >15 axes studied in serial sections, we describe a new anatomically preserved rhyniopsid from the Battery Point Formation, *Eddianna gaspiana* gen. & sp. nov.. *Eddianna* axes are up to 2 mm in diameter and have a well-developed terete xylem strand with potential centrarch maturation (comprising 80% of the cross sectional surface area) that features *Sennicaulis*-type tracheid wall thickenings. A thin layer interpreted as phloem is preserved around the central xylem and an irregular sclerenchymatous cortex forms longitudinal anastomosing ridges on the outside of the axes. The anatomy of *Eddianna* axes suggests that they represent lower portions, specialized in efficient water transfer, of a larger plant whose distal regions have yet to be discovered. *Eddianna*, the first permineralized rhyniopsid described from the Battery Point Formation, is one of only four anatomically preserved plants reported from this unit. These fossils reiterate the potential for additional discoveries of anatomically preserved plants in the Battery Point Formation.

## INTRODUCTION

Vascular plants underwent their first major phase of evolutionary radiation in the Early Devonian (Banks, 1970; Gensel & Andrews, 1984; Kenrick & Crane, 1997; Gensel & Edwards, 2001; Gensel, 2008; Cascales-Miñana *et al.*, 2010; Hao & Xue, 2013). Rocks from this time interval preserve some of the oldest vascular plants, which provide information about the evolution of vascular tissues and tracheophyte diversity. While numerous plant species have been documented from Lower Devonian strata, many of these species are incompletely understood. This underscores a need for continued investigations of Early Devonian floras. The majority of Early Devonian plants had simple organization, consisting of undifferentiated, branched photosynthetic axes. These plants have been traditionally assigned to a few main groups, one of which is the Rhyniopsida (Kenrick & Crane, 1997). The morphological simplicity of rhyniopsids and other Early Devonian plants, in general, provides only a dearth of characters for comparisons, hindering studies of both the systematics and relationships of these plants. In this context, anatomical information on Early Devonian plants can contribute crucial resolution to these questions and discovery of new anatomically preserved fossils continues to be a primary goal of paleobotanical investigations.

The Rhyniaceae, including *Rhynia* Kidston & Lang (1917), *Stockmansella* Fairon-_Demaret (1986), and *Huvenia* Hass & Remy (1991), were recognized as a group by Hass and Remy (1991). Phylogenetic analyses by Kenrick & Crane (1991; 1997) recovered this grouping as a clade, renamed Rhyniopsida (Kenrick & Crane, 1997), that shared a distinctive type of adventitious branching, sporangia attached to a specialized pad of tissue, and an abscission or isolation layer at the base of the sporangium. Kenrick & Crane (1991, 1997) noted that the three genera share the same type of water-conducting cells seen in *Sennicaulis* Edwards (1981), termed S-type tracheids, which indicates that the latter is also a member of the Rhyniopsida and the S-type tracheids are another defining feature of this group. Gerrienne *et al.* (2006) have proposed the phylum (division) Paratracheophyta for the group that includes rhyniopsids (*Rhynia*, *Stockmansella*, *Huvenia*, *Sennicaulis*) along with *Taeniocrada dubia* Kräusel & Weyland and the gametophytes *Remyophyton* Kerp, Trewin & Hass and *Sciadophyton* Steinmann.

*Rhynia gwynne-vaughanii* Kidston & Lang (1917) is known from the Pragian Rhynie chert as silica permineralizations (Edwards, 1980). *Sennicaulis hippocrepiformis* Edwards (1981) has been described from pyrite-limonite permineralizations in the Pragian Senni Beds of the Old Red Sandstone, in Wales. *Stockmansella*, with two species – *S. langii* Fairon-Demaret, (1985; 1986) and *S. remyi* Schultka & Hass (1997) – was described from compressions that preserve pyrite tracheid casts in the Pragian Bois d’Ausse Layers of Belgium and the Eifelian Brandenberg Formation of Germany. *Huvenia* also includes two species – *H. kleui* Hass & Remy (1991) and *H. elongata* Schultka (1991) – reported in the Pragian Wahnbach Schichten and the Emsian Nellenkopfschichten of Germany, and preserved as compressions containing pyrite tracheid casts.

In North America, rhyniopsids have been reported from the Battery Point Formation (Hotton *et al.*, 2001; Gensel, 2005), as part of a very diverse Early Devonian flora. The Battery Point Formation represents sediments deposited in braided fluvial to coastal environments (Cant and Walker, 1976; Griffing, Bridge & Hotton, 2000). These Emsian rocks (McGregor, 1977) host plants preserved as carbonaceous compressions and anatomically preserved permineralizations. Compressed fossils, known since John William Dawson’s early explorations (e.g., Dawson, 1859), have been extensively characterized (e.g., Gensel & Andrews 1984; Hotton *et al*., 2001 and, more recently, Gensel & Berry, 2016), and make this the most diverse Early Devonian flora in North America. In contrast, permineralized specimens of the Battery Point Formation plants have not received as much attention. Only a handful of studies have addressed anatomically preserved fossils of this unit (Banks & Davis, 1969; Banks, Leclercq & Hueber, 1975; Banks, 1981; Hartman & Banks, 1980; Hartman, 1981; Banks & Colthart 1993; Hoffman & Tomescu, 2013), which continues to yield new permineralized material. Here, we describe a new rhyniopsid genus from anatomically preserved material in the Battery Point Formation. This study is part of renewed efforts aimed at fully exploring the diversity of the permineralized flora of this important Early Devonian unit.

## MATERIAL AND METHODS

Eighteen axes of this new fossil are preserved by calcareous cellular permineralization in four cobbles collected by Dr. Francis M. Hueber (Smithsonian Institution—NMNH) in 1965, from the Battery Point Formation, in the vicinity of Douglastown. This unit is exposed between Douglastown and Tar Point, on the southern shore of Gaspé Bay, Québec, Canada, and its age increases from late Emsian at Douglastown to early Emsian at Tar Point (McGregor, 1977). Therefore, the age of the fossils described here is mid-to late Emsian, ca.402–394 million years old (Cohen *et al*., 2016). Hosting a broad diversity of mid-to late Emsian plants, the cobbles containing this new fossil are composed of sediments deposited in braided fluvial to costal environments (Cant & Walker, 1976; Griffing *et al*., 2000).

The fossils were studied in serial anatomical sections obtained using the cellulose acetate peel technique (Joy, Willis & Lacey, 1956). Slides for bright-field microscopy were mounted with Eukitt (O. Kindler, Freiburg, Germany) mounting medium. Images were captured using a Nikon Coolpix 8800VR digital camera mounted on a Nikon E400 compound microscope. Material for scanning electron microscopy was obtained from cellulose acetate peels using the method detailed in Matsunaga *et al*. (2013). SEM images were generated using a FEI Quanta 250 (Hillsboro, Oregon, USA). Images were processed using Adobe Photoshop (San Jose, California, USA). All cobble slabs, acetate peels and slides are stored in the U.S. National Museum of Natural History – Smithsonian Institution (USNM no. 557783-5, 557840, 557790-1, 557790-3).

## TERMINOLOGY

Several authors (Kenrick & Crane, 1991; Kenrick, Edwards & Dales, 1991; Edwards *et al.*, 2003; Gerrienne *et al.*, 2006) have discussed the possibility that S-type conducting cells followed a developmental pathway different from those seen in the tracheids of extant vascular plants. This is contrary to the view of a single origin of tracheids, including S-type conducting cells, in land plants (Cook & Friedman, 1998). The data we present here do not contribute to this debate in either direction. While acknowledging that it is unclear whether S-type conducting cells are homologous to tracheids, for simplicity we are referring to them as tracheids and to the tissue they form as xylem. For the same reason, we use the term protoxylem to refer to the area comprising the smallest water-conducting cells, and metaxylem for the rest of the water-conducting cells, which are larger, even though developmental evidence for the timing of their maturation with respect to tissue elongation is lacking.

## SYSTEMATICS

### Plesion Rhyniopsida Kenrick & Crane

#### ***Eddianna*** Pfeiler & Tomescu **GEN. NOV.**

*Diagnosis*: Axes with large amount of xylem and scant extraxylary tissues including phloem and cortex. Cortex sclerenchymatous, irregular, forming anastomosing longitudinal ridges. Central xylem strand terete, with potential centrarch maturation. Tracheids with annular to helical *Sennicaulis*-type wall thickenings.

***Derivation****: Eddianna* is named in honor of Prof. Dianne Edwards, in recognition of her significant contributions to Silurian and Devonian palaeobotany.

***Type species****: Eddianna gaspiana*

#### ***Eddianna Gaspiana*** Pfeiler & Tomescu **SP. NOV.**

***Diagnosis***: Axes up to 2 mm or more in diameter, with central xylem strand occupying ca. 80% of cross-sectional surface area. Cortex sclerenchymatous, at least three cells thick, with cells up to 48 µm in diameter. Cortex forms longitudinal ridges up to ca. 190 µm tall and 480 µm across at the base. Phloem thin, 48-72 µm, consisting of 1-3 layers of narrow elongated cells with thin, straight walls. Central xylem strand circular-elliptical in cross section, up to 1.6 mm or more in diameter. Metaxylem tracheids round to oval in cross section, up to 40-66 µm in diameter. Tracheid wall thickenings with spongy structure and lined with a microporate layer. Wall thickenings ca. 12 µm wide, spaced at ca. 24 µm apart, and protrude ca. 7 µm into the cell lumen. Pore density in microporate layer ca. 16 µm^-2^; pore diameter 0.15-0.20 µm. Larger axes with concentric zonation of the metaxylem: larger tracheids in the outer zone and narrower tracheids in the inner zone; limit between the two zones marked by discontinuous ring of even narrower tracheids.

***Derivation of name****: gaspiana* for the Gaspé Peninsula (Canada) where the specimens were collected.

***Holotype***: axis in slabs USNM 557840 Gbot, Htop.

***Paratypes***: axes in slabs USNM 557840 Hbot, 557783-5 Atop, 557790-1 Bbot-Ctop, 557790-3 Atop.

***Locality and stratigraphy:*** Battery Point Formation, in the vicinity of Douglastown; mid-to late Emsian, ca.402–394 Ma.

## DESCRIPTION

The axes of *E. gaspiana* range in diameter from 0.6 mm to 2.1mm. The longest observed fragment measures 2.5 cm. Reproductive structures have not yet been found associated with any of the axes. External evidence for axis branching is also missing, although one specimen shows anatomical evidence for branching.

Specimen 557840 Gbot-Htop is an axis cut in cross section (Fig. 1A). The axis is 1.25 mm in diameter. The cortex consists of highly sclerified cells that form a dark layer in which individual cells are difficult to discern. The thickness of the cortex is variable, including irregular protrusions that range up to 144 µm tall and 192 µm across the base, as well as regions where the cortex can be very thin (12 µm), possibly as a result of incomplete preservation. Beneath the cortex is a thin layer 48 µm wide. Cellular preservation in this layer is incomplete, but fragments of cell layers indicate that it consisted of parenchyma cells. Based on its position immediately outside the xylem cylinder, as well as its parenchymatous nature, we interpret this layer as a photosynthate-conducting tissue and refer to it as phloem, while acknowledging that its homology with tracheophyte phloem is equivocal. The central xylem strand is circular in shape, 1.09 mm in diameter. Although cellular detail is blurred toward the center of the xylem, the centripetal increase in tissue density (Fig. 1A) is consistent with a centrarch pattern of maturation and is, thus, tentatively interpreted as such. Cells walls of the xylem tracheids in this and all other specimens have a conspicuously lighter color compared to the xylem of all other plants in the assemblage. Tracheids have round to oval outlines with diameters up to 30-42 µm.

**Figure 1.**
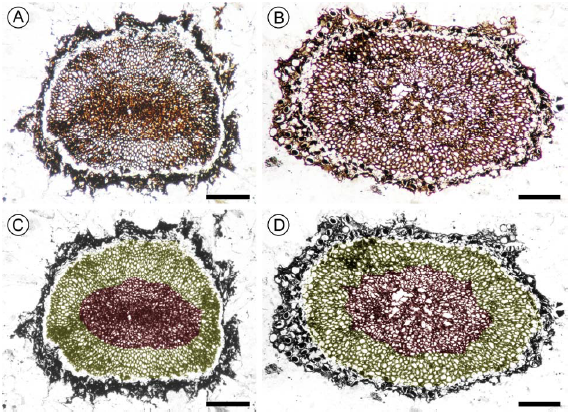
Eddianna gaspiana gen. & sp. nov. A, B, Axis cross sections exhibiting large primary xylem with potential centrarch maturation cylinders separated from thin, irregular sclerenchymatous cortex by a thin layer of phloem; note incomplete preservation of xylem at center in B. C, D, same images as in A and B, respectively, colorized to emphasize concentric zonation of the metaxylem, with larger tracheids in the outer zone (yellow) and narrower tracheids in the inner zone (brown); a thin discontinuous layer of very fine tracheids (more conspicuous in B, D) marks the border between the two zones; note subtle radial patterning of tracheids, locally, in the outer zone, and pattern of decreasing tracheid diameter toward xylem periphery. A, C, USNM-557840 Htop #20a; B, D, USNM-557790-1 Bbot #88a. Scale bars =300 µm.

Specimen 557790-1 Bbot-Ctop is another axis cut in cross section, 2.1 mm in diameter (Fig. 1B). The cortical cells, easily discernible, are 24-48 µm in diameter and have sclerified walls (Fig. 4A). Cortical protrusions are up to 192 µm tall and 480 µm across at the base. The phloem layer is 50-70 µm wide, and is composed of 1-3 layers of well-preserved thin-walled cells (Fig. 4A). The xylem cylinder is 1.58 mm diameter and metaxylem tracheids are up to 42-66 µm in diameter. The potential centrarch pattern of xylem maturation is less conspicuous in this specimen, due to the presence of several voids in the central area of the xylem (Fig. 1B); it is unclear if these areas of missing tissue are the result of taphonomy or development.

Specimen 557840 Hbot exhibits two axes cut in longitudinal section (Fig. 2A). Both axes are at least 2.5 cm in length. In these longitudinal sections, the cortex shows important variations in thickness, as well as numerous discontinuities (Fig. 2B). The thickness of the cortex is up to 1.2 mm and individual cells are difficult to discern. The phloem layer is up to 72 µm and 5-6 cells thick, and consists of narrow cells with thin, straight walls (Fig. 3A, B). These cells are elongate, but we did not observe end walls so their length cannot be measured. In a radial longitudinal section the xylem strand is 0.8 mm in diameter and exhibits more conspicuous centrarch maturation (Fig. 3C). Like in the phloem, the length of the tracheids cannot be measured due to the lack of end walls. Tracheids are up to 48 µm in diameter and have annular to helical thickenings. The thickenings are ca. 12 µm wide and spaced at ca. 24 µm apart. They protrude ca. 7 µm into the cell lumen. Scanning electron microscopy revealed the spongy nature of the tracheid wall thickenings (Fig. 3D), which are lined with a microporate inner layer (pore density is ca. 16 µm^-2^ and pore diameter ca. 0.15-0.20 µm) (Fig. 4B). Both these features are characteristic of *Sennicaulis*-type (S-type) tracheids (Kenrick *et al.*,1991a).

**Figure 2.**
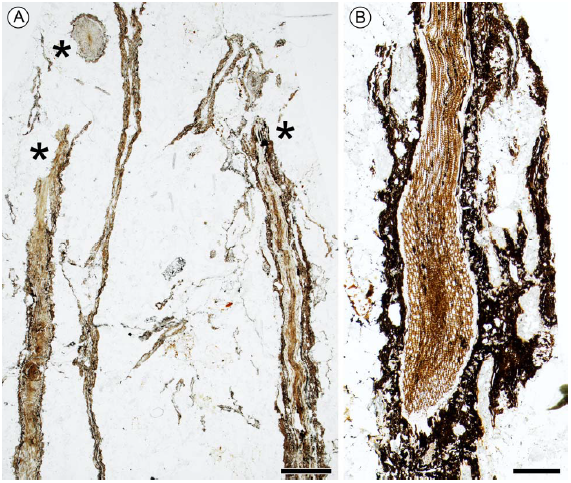
*Eddianna gaspiana* gen. & sp. nov. A, Three ***Eddianna*** axes (asterisks); one is sectioned transversally and the other two longitudinally; the other plant material is ***Psilophyton***. USNM-557840 Hbot #117f. Scale bar = 2 mm. B, Oblique longitudinal section of axis with prominent xylem strand (light brown) and dark, sclerenchymatous cortex forming anastomosing longitudinal ridges (very pronounced on the right). USNM-557840 #1f. Scale bar = 600 µm.

**Figure 3.**
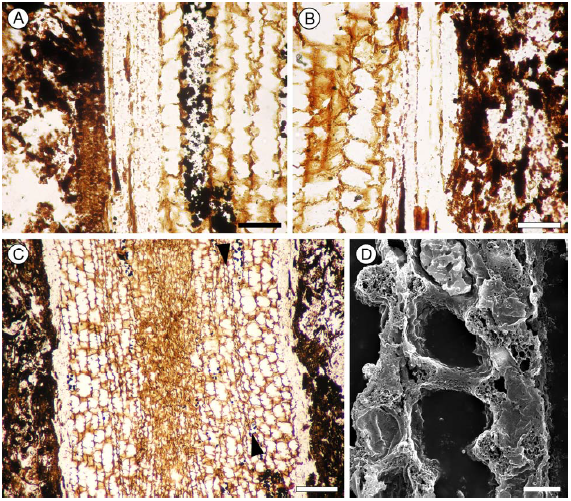
*Eddianna gaspiana* gen. & sp. nov. A, B, Longitudinal sections of axis showing (right to left in A and left to right in B) metaxylem tracheids (light brown) with conspicuous helical wall thickenings; phloem layer (light brown) with narrow cells and fine, straight vertical cell walls; and incompletely preserved sclerenchymatous cortex (dark brown). Scale bars = 50 µm. C, Longitudinal radial section of axis with large xylem strand (protoxylem compressed, distorted at center); sclerenchymatous cortex (dark brown, at left and right); and thin phloem sandwiched between xylem and cortex; note conspicuous helical thickenings of metaxylem tracheids and fine band of much narrower tracheids (between arrowheads) between the two concentric zones of the metaxylem (e.g. Fig. 1B, 1D). Scale bar = 150 µm. USNM-557840 Hbot #1f. D, Scanning electron micrograph of tracheids with helical wall thickenings, note spongy structure of the thickenings, characteristic of *Sennicaulis*-type tracheids. USNM-557840 Hbot #44. Scale bar = 10 µm.

**Figure 4.**
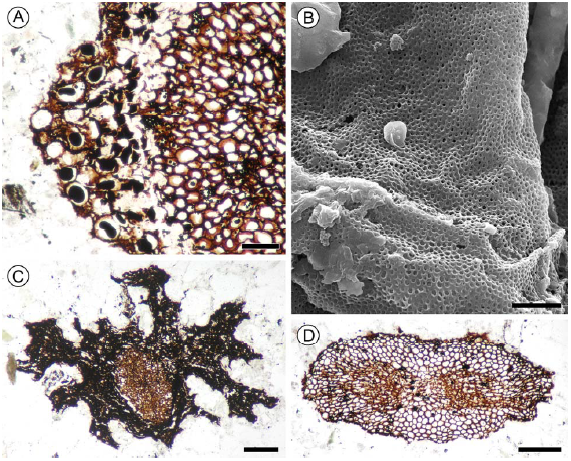
*Eddianna gaspiana* gen. & sp. nov. A, Cross section of axis; here the cortex (left) has larger cells with thinner walls, some filled with dark content; the 2-3 cells-thick phloem is somewhat compressed between the cortex and metaxylem; the metaxylem (right) shows robust helical wall thickenings. USNM-557790-1 Bbot #88a. Scale bar = 100 µm. B, Scanning electron micrograph of the microporate inner lining of the *Sennicaulis*-type tracheid wall. USNM-557840 Hbot #44. Scale bar = 2 µm. C, Cross section of smaller axis with well-developed sclerenchymatous cortex; note numerous robust irregular protrusions of the cortex, which form anastomosing longitudinal ridges along axes. USNM-557783-5 Atop #8a. Scale bar = 200 µm. D, Cross section of decorticated axis at branching point, exhibiting elliptical xylem with two protoxylem strands. USNM-557790-3 Atop #30a. Scale bar = 200 µm.

Specimen 557783-5 Atop is a smaller axis cut in cross section (Fig. 4C). This axis is 1.2 mm in diameter, with a xylem strand 480 µm in diameter. Cellular detail of both the xylem and cortex is not clear as they are both very dense. This specimen is interesting in exhibiting very prominent, irregular cortical protrusions. The protrusions are elongated radially and have blunt or pointed tips; some are branched. Together with the longitudinal sections in 557840 Hbot, this indicates that the sclerenchymatous cortex formed vertical ridges that branched and anastomosed irregularly.

Specimen 557790-3 Atop is a decorticated axis cut in cross section. The xylem has a flattened, oval outline and two areas of protoxylem surrounded by smaller metaxylem tracheids (Fig. 4D). This anatomy reflects branching, which is likely isotomous.

The majority of specimens show subtle yet distinct concentric zonation of the metaxylem (Fig. 1C, D). In the outer zone, tracheids are larger, whereas in the inner zone they are narrower. The limit between the two zones is relatively sharp and marked by a discontinuous ring of even narrower tracheids (Figs. 1B, 3C). Tracheid diameter also decreases slightly toward the periphery of the xylem cylinder (Fig. 1A, B). Tracheids in the outer zone exhibit a subtle radial patterning in shape and arrangement in some areas of the cross sections (Fig. 1A, B).

## JUSTIFICATION FOR A NEW GENUS

According to Kenrick & Crane (1997), the Rhyniopsida are early tracheophytes with simple organography characterized by adventitious branching of axes and by sporangia that are attached to a specialized pad of tissue and have an abscission layer at the base. Additionally, all four genera of known rhyniopsids – *Rhynia*, *Sennicaulis*, *Stockmansella*, and *Huvenia* – have tracheids characterized by S-type wall thickenings (Kenrick & Crane, 1997; Kenrick *et al*., 1991a; Kenrick & Crane, 1991; Kenrick, Remy & Crane, 1991b). The *Eddianna* material described here is strictly vegetative and does not provide evidence for adventitious branching. Nevertheless, the presence of S-type tracheid thickenings places this plant among the rhyniopsids, the only group that exhibits this type of tracheids.

Compared to the other rhyniopsid genera (Table 1), *Eddianna* stands out by its large amount of xylem with respect to the axis diameter and the proportion of extraxylary tissues. Whereas in the other genera the xylem occupies<10% of the cross-sectional surface area of axes, in *Eddianna* the xylem occupies >80%. On the other hand, while the other rhyniopsids have a thick parenchymatous cortex, *Eddianna* has very little extraxylary tissues. Additionally, the axes of *Eddianna* ssare covered by pronounced longitudinal ridges that branch and anastomose, unlike any of the other rhyniopsids. Finally, it is worth noting that the *Eddianna* has the largest tracheids of all these plants. Taken together, these differences imply that *Eddianna* is different form all previously described rhyniopsids, thus warranting erection of a new genus.

**Table 1.** Rhyniopsid genera compared.

|  | *Eddianna* | *Stockmansella* | *Huvenia* | *Sennicaulis* | *Rhynia* |
| --- | --- | --- | --- | --- | --- |
| Max. axis diameter | 2.1 mm | 15.0 mm | 15.0 mm | 6.0 mm | 2.0 mm |
| % Xylem surface | 82.0 | 0.7 - 2.6 | 1.6 - 6.3 | 2.3 | 0.5 |
| Max. tracheid lumen diameter | 67 µm | 50 µm | 27 µm | 45 µm | 21 µm |
| Tracheid wall thickening width | 6.0 - 21.6 µm | 4.7 - 9.5 µm | 3.5 - 6.3 µm | 6.3 - 27.5 µm | 10.7 µm |
| Extraxylary tissues | overall thin; thin-walled phloem cells, sclerenchymatous cortex | thick cortex ?parenchymatous | thick cortex ?parenchymatous | thick cortex ?parenchymatous thick-walled cells in outermost layers | thick cortex parenchymatous |
| Projections on axis surface | low cones; irregular anastomosing ridges | hemispherical rhizoid-bearing bulges | longitudinally ovoid low relief | ?smooth | hemispherical rhizoid-bearing bulges |
| Age | Emsian | Pragian - Eifelian | Pragian - Emsian | Pragian | Pragian |
| Location | Québec, Canada | Belgium, Germany | Germany | Wales | Scotland |
| Preservation | carbonate permineralization | compression, tracheid casts | compression, tracheid casts | pyrite-limonite permineralization | silica permineralization |
| References | this study | Fairon-Demaret 1985; Schultka & Hass 1997 | Hass & Remy 1991; Shultka 1991; Kenrick & Crane 1991; Schultka & Hass 1997 | Edwards 1981; Kenrick *et al.* 1991 | Kidston & Lang 1917; Edwards 1980 |

All previously known rhyniopsid genera have thick extraxylary tissues. This raises the question of incomplete preservation: could *Eddianna* represent specimens of a plant with thick parenchymatous extraxylary tissues [such as the compressions referred to as *Taeniocrada dubia* by Hueber (1982) and “*Taeniocrada dubia*“/new genus D by Hotton *et al.* (2001)] that were not preserved? This possibility needs to be considered, since the permineralized plant assemblage that hosts *Eddianna* is allochthonous, therefor?e the plant material underwent transport which may have affected its integrity. Additionally, an epidermal layer cannot be distinguished on the *Eddianna* axes. This could be explained either by sclerification of the epidermis along with the underlying cortical layers or by loss, which would indicate incompleteness of preservation of extraxylary tissues.

Of the relatively numerous (18) specimens studied, none shows evidence of tissues, even incomplete, outside the sclerenchymatous cortex. Instead, fragments of other plants are found alongside and in contact with the sclerenchymatous cortex of *Eddianna* axes. On one hand, these are consistent with an interpretation of the *Eddianna* specimens as being anatomically complete as described here. On the other hand, there observations do not unequivocally reject the hypothesis of taphonomic loss of a putative parenchymatous outer cortex. However, it is important to note that the multiple *Eddianna* specimens are found in four separate rock samples that exhibit differences in grain size (from silt-size to microconglomerate). This observation has two implications: (i) the specimens come from different populations, thus they have undergone transport over different distances; and (ii) they underwent transport under different flow regimes. Consequently, our sample of *Eddianna* specimens covers a whole range of intensities of exposure to taphonomic factors. This makes it likely that at least fragments of an outer parenchymatous cortex, if such a layer was present, would have been preserved in at least some of the specimens. Because we observed no such structures, the interpretation that *Eddianna* had an outer parenchymatous cortex that was not preserved, is not supported by any physical evidence. Instead, the weight of all evidence associated with the *Eddianna* specimens supports the hypothesis that an outer parenchymatous cortex was not present and the specimens preserve relatively complete anatomy of the axes, which we adopt as our interpretation of these fossils.

## FUNCTIONAL ANATOMY AND THE *EDDIANNA* PLANT

The thick sclerified cortex of *Eddianna* could have had a role in providing both mechanical support for the axes and protection from desiccation and herbivory (i.e. piercing-and-sucking; Labandeira & Currano, 2013). The primarily sclerenchymatous nature of the cortex also indicates that this layer had little to no photosynthetic capabilities. Additionally, the only other extraxylary tissue of the *Eddianna* axes is a thin phloem layer. Together, these suggest that *Eddianna* axes were not photosynthetic. Concurrently, the axes have a high proportion of xylem which is consistent with high water transport capacity. Together, the lack of photosynthetic tissues and the high water transfer capacity of the axes suggest that they represent the lower portions of larger plants that conducted photosynthesis in more distal parts of hypothetical branching systems. Therefore, the lower portions of these plants consisted of stiff, nonphotosynthetic axes built for maximizing support and water transport.

Basal axes that have almost exclusively a water conduction role, such as those of *Eddianna*, are justified only if (1) water is readily available in the substrate; (2) the plants have an extensive absorptive rooting system or (3) they have extensively developed distal photosynthetic parts; or a combination of these conditions. Therefore, it is likely that the *Eddianna* plant grew in a water-rich environment, e.g. in the immediate vicinity of water bodies, on floodplains, or on substrates with a high water table. Alternatively, the distal photosynthetic parts of *Eddianna* could have been extensively developed (highly branched), or had rapid growth, or maintained high turgidity, all of which require high water intake. It is also worth considering that rhyniopsids have adventitious branching (Kidston & Lang, 1917; Kenrick & Crane, 1997; Kearney, Kerp & Hass, 2016) wherein lateral axes, attached to the plant by constricted bases marked by a discontinuous vascular strand, could be abscised and act as propagules (Kearney *et al*., 2016). This rhyniopsid feature suggests yet another possible explanation for the high xylem volume of *Eddianna* axes: high turnover of distal photosynthetic parts by abscission of such adventitious deciduous branches, which would have also required high water use.

The concentric zonation observed in the xylem of *Eddianna* axes is puzzling. Although weak/faint radial patterning is present in some xylem sectors of these axes, this does not provide evidence for secondary growth. Therefore, the concentric zonation cannot be attributed to variations in growth rhythm of a lateral meristem (cambium). However, this concentric pattern is consistently present in many *Eddianna* specimens, which indicates that it reflects a constitutive feature of development in this plant.

## RHYNIOPSIDS IN THE BATTERY POINT FORMATION FLORA

Because ***Eddianna*** is a rhyniopsid, a logical question is: are these fossils part of another, previously described, rhyniopsid from the Battery Point Formation? Rhyniopsids reported previously from the Battery Point Formation include only compression fossils: *Huvenia* sp. nov. (Hotton *et al*., 2001), a type referred to as ‘new genus D’ or “*Taeniocrada dubia*” (Hotton *et al*., 2001 and Hueber 1982), and fossils referred to as *Stockmansella* (Hotton *et al*., 2001) or cf. *Stockmansella* (Gensel, 2005). *Huvenia* sp. nov. axes are about 5 mm in diameter and the axes of new genus D (“*Taeniocrada dubia*“) are 9-20 mm in diameter (Hotton *et al*., 2001). The sizes of cf. *Stockmansella* axes in the Battery Point Formation are not reported, but the axes of the two *Stockmansella* species described to date are 2-15 mm in diameter (Fairon-Demaret, 1985; Schultka & Hass, 1997). With the exception of the smallest *Stockmansella* specimens, all of these plants have axes that are much wider than those of *Eddianna*. Structurally speaking, in most plants, growth and branching patterns lead to progressively thinner axes distally. Therefore, if ***Eddianna*** axes represent the lower portions of the plant, the distal parts of this plant are predicted to consist of axes of smaller diameter than the lower portions. Consequently, the distal parts of the *Eddianna* plant would be much thinner than the axes of the rhyniopsids previously described from the Battery Point Fomation. This excludes these other rhyniopsids as potential candidates for the distal parts of *Eddianna*, the search for which is ongoing.

## CONCLUSIONS

We are describing a new anatomically preserved plant from Emsian deposits of the Battery Point Formation (Gaspé Canada). The plant, ***Eddianna gaspiana*** gen. et sp. nov., has distinctive anatomy characterized by a thick xylem strand with S-type tracheids and a ridged sclerenchymatous cortex, which set it apart from previously described rhyniopsids. The anatomy of ***Eddianna*** fossils indicates that they represent the lower portions of a plant whose distal regions have yet to be discovered. ***Eddianna*** is the first anatomically preserved rhyniopsid reported from the Battery Point Formation. This unit has already yielded compression fossils potentially representing three types of rhyniopsids, but the size of these rhyniopsid axes is inconsistent with predicted sizes of the distal parts of *Eddianna*.

***Eddianna*** adds a fourth type to the diversity of plants known from anatomically preserved specimens in the Battery Point Formation. Previously described permineralized fossils represent trimerophytes – *Psilophyton* (Banks *et al*., 1975), *Franhueberia* (Hoffman & Tomescu, 2013) and a third unnamed type (Gensel, 1984) – and a zosterophyll (*Crenaticaulis*; Banks & Davis, 1969). In contrast, the compressions in the Battery Point Formation represent a higher diversity of plants (Gensel & Andrews, 1984; Hotton *et al*., 2001; Gensel, 2005). This higher diversity should be mirrored in the permineralized component of the flora. Together, these suggest that the Battery Point Formation hosts additional diversity of permineralized species that await in-depth investigation. This study is part of renewed efforts to characterize this diversity.

## ACKNOWLEDGEMENTS

We thank Francis M. Hueber, who collected the specimens containing ***Eddianna*** axes. We are indebted to William DiMichele, Carol Hotton and Jonathan Wingerath (National Museum of Natural History – Smithsonian Institution) for facilitating specimen loans; Kelly K.S. Matsunaga (University of Michigan) for help during work in the NMNH collections; and Alejandro C. Bippus, Selin Toledo, Marty Reed and Casey R. Lu for assistance with scanning electron microscopy. The research has been supported by a grant (to AMFT) from the American Philosophical Society, which is gratefully acknowledged.

